## SupplementaryMaterial for "Aberrant integration of Hepatitis B virus DNA promotes major restructuring of human hepatocellular carcinoma genome architecture"

**SUPPLEMENTARY FIGURES**

#### Supplementary Figure 1

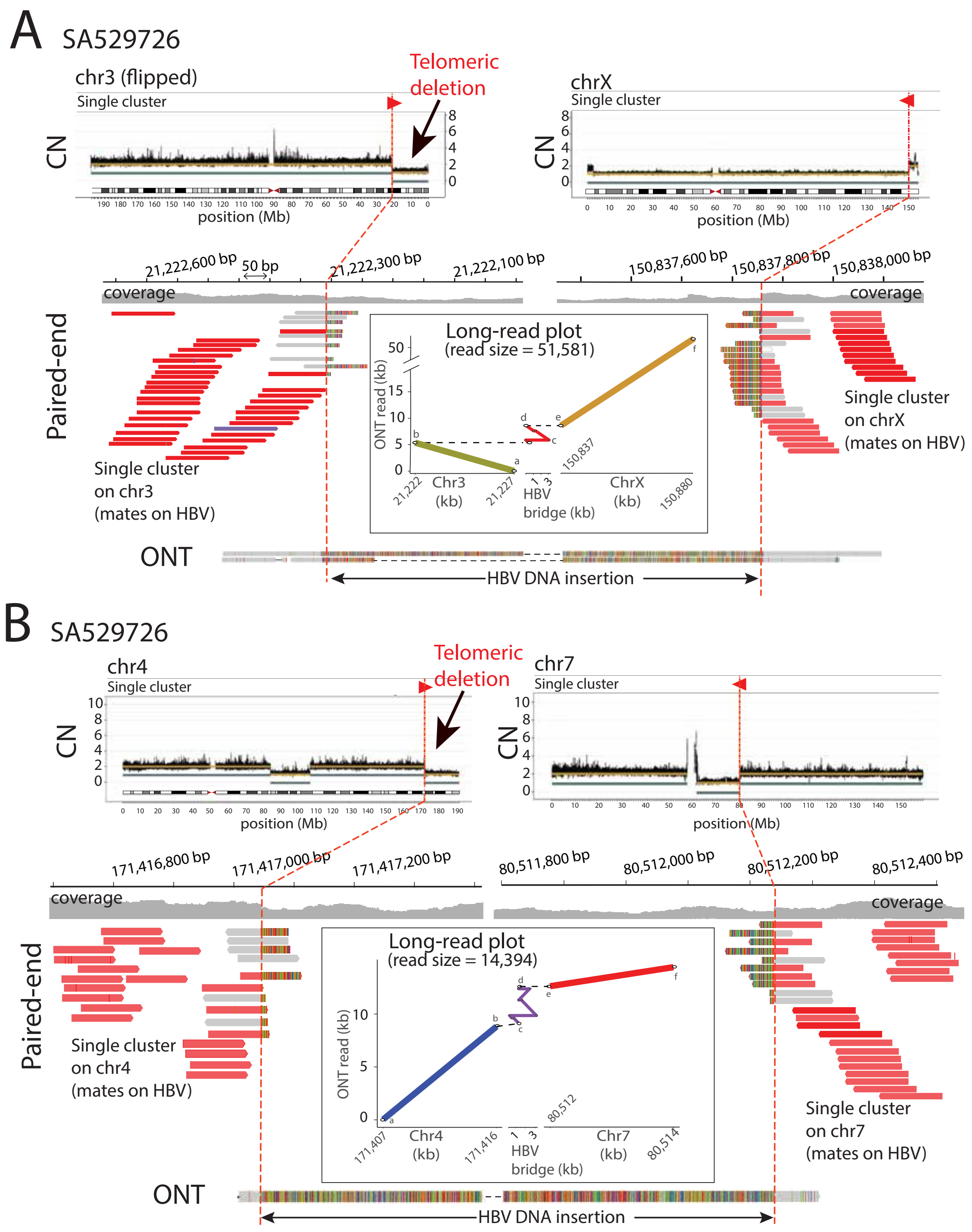


**Supplementary Figure 1. HBV-mediated interchromosomal rearrangements revealed by ONT sequencing in HCC tumour SA529726. (a)** The Illumina paired-end sequencing data (short-reads in red) shows one positive cluster of discordant read-pairs pointing to an HBV insertion and demarcating the boundary of a 21.2 Mb telomeric deletion on 3p (note the orientation of the chromosome is inverted for illustrative purposes). The copy number plot (CN) at the top shows the total (gold line) and minor (grey line) chromosomes' copy number profiles. The ONT data at the bottom shows two long-reads obtained with Oxford Nanopore that reveals one 3,268 bp long HBV DNA insertion bridging the interchromosomal rearrangement between 3p and Xq. A negative cluster of Illumina reads confirms the presence of the other extreme of the HBV insertion on Xq. The long-read plot represents the alignment of one ONT long-read to chromosomes 3 and X of the human reference genome and an HBV consensus sequence, which validates the interchromosomal rearrangement mediated by the virus. Note that the HBV insertion shows a typical rearranged pattern. **(b)** In the same sample, an HBV-mediated interchromosomal rearrangement between chromosomes 4 and 7 removes 19.8 Mb on 4q, including the telomere. Two paired-end clusters (short-reads in red) point to an HBV event. One ONT long-read reveals a 3,494 bp long HBV DNA insertion that bridges the translocation between 4q and 7q (long-read plot).

### Supplementary Figure 2

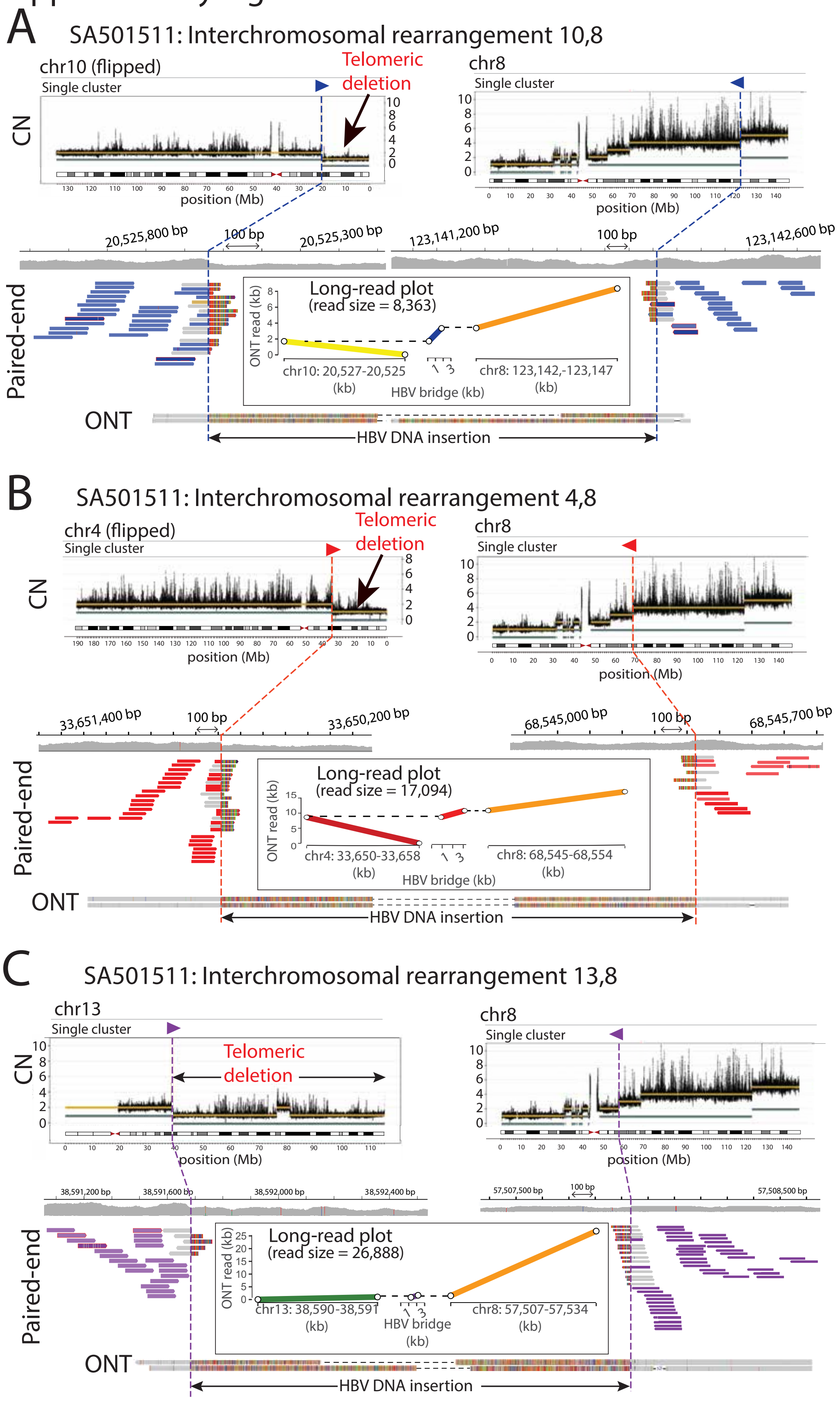

**Supplementary Figure 2. HBV-mediated interchromosomal rearrangements revealed by ONT sequencing in HCC tumour SA501511. (a)** The Illumina paired-end data (short-reads in blue) shows two clusters, one on chromosome 10 and another on chromosome 8, which point to both extremes of an HBV insertion. The copy number (CN) plot at the top shows the total (gold line) and minor (grey line) chromosomes' copy number profiles, revealing a 20.5 Mb telomeric deletion on 10p associated with the HBV insertion event (note that the CN plot on chromosome 10 is flipped for illustrative purposes). The ONT data at the bottom shows two long-reads that reveal the real configuration of the rearrangement. In the long-read plot, a 1,629 bp HBV insertion bridges an interchromosomal rearrangement between 10p and 8q. **(b)** The Illumina paired-end data (short-reads in red) shows two clusters, one on chromosome 4 and another on chromosome 8, which point to both extremes of an HBV insertion. The CN plot at the top reveals a 33.6 Mb telomeric deletion on 4p associated with the HBV insertion event (note that the CN plot on chromosome 4 is flipped for illustrative purposes). The ONT data at the bottom shows two long-reads that reveal the real configuration of the rearrangement. In the long-read plot, a 2,227 bp HBV insertion bridges an interchromosomal rearrangement between 4p and 8q. **(c)** The Illumina paired-end data (short-reads in purple) shows two clusters, one on chromosome 13 and another on chromosome 8, which point to both extremes of an HBV insertion. The CN plot at the top reveals a 76.7 Mb deletion on 13q involving the telomere, which is associated with the HBV insertion event. The ONT data at the bottom shows two long-reads that reveal the real configuration of the rearrangement. In the long-read plot, a 531 bp HBV insertion that bridges an interchromosomal rearrangement between 13q and 8q.

### Supplementary Figure 3

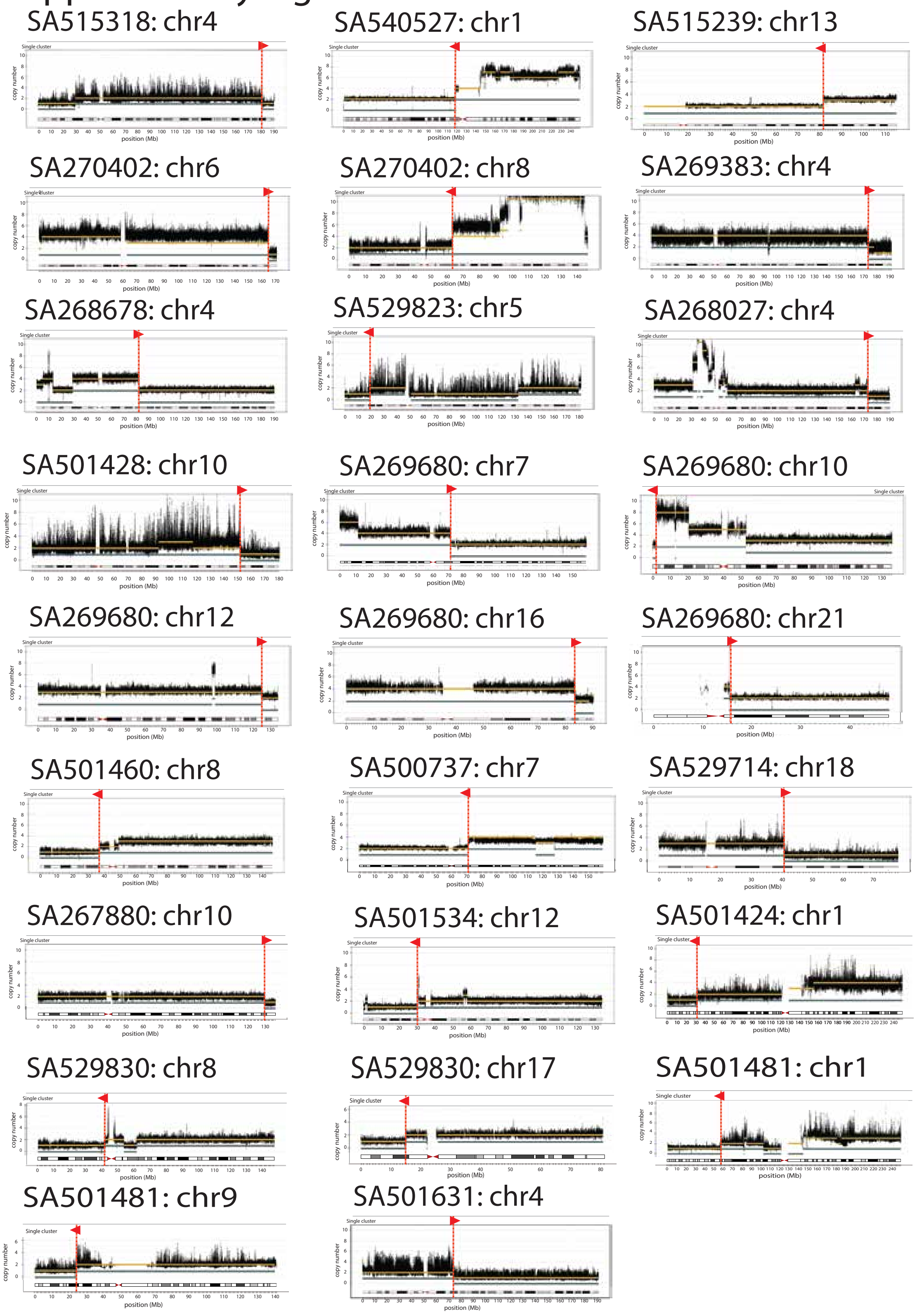


**Supplementary Figure 3. Telomeric-like deletions associated with HBV DNA insertions are frequent in human HCC.** Copy number (CN) plots from 26

chromosomes with telomeric-like deletions associated with HBV insertion events in HCC tumours from the PCAWG dataset. The CN plots show the total (gold line) and minor (grey line) chromosomes' copy number profiles. Arrows indicate the orientation (positive or negative) of the paired-end clusters supporting each HBV insertion.

### Supplementary Figure 4

## SA529830

Tumour Purity: 0.429

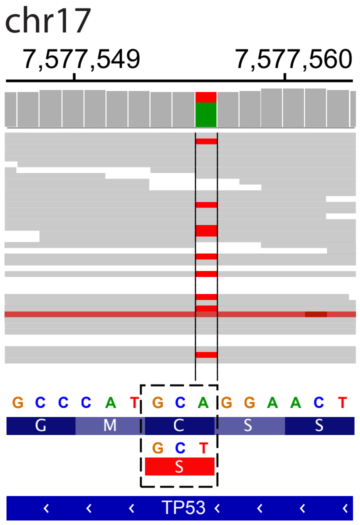


**Supplementary Figure 4. Inactivating mutation in *TP53* from tumour SA529830.** Inactivating mutation detected in the second allele of *TP53* from sample SA529830. The picture corresponds to an Integrative Genomics Browser (IGV) view of the paired-end mapping data at the relevant locus. An A>T mutation at exon 7 of the gene involves the aminoacid change C242S, catalogued as loss-of-function mutation [23]. Note that tumour purity is 0.429, which explains the reference 'A'.

Supplementary Figure 5

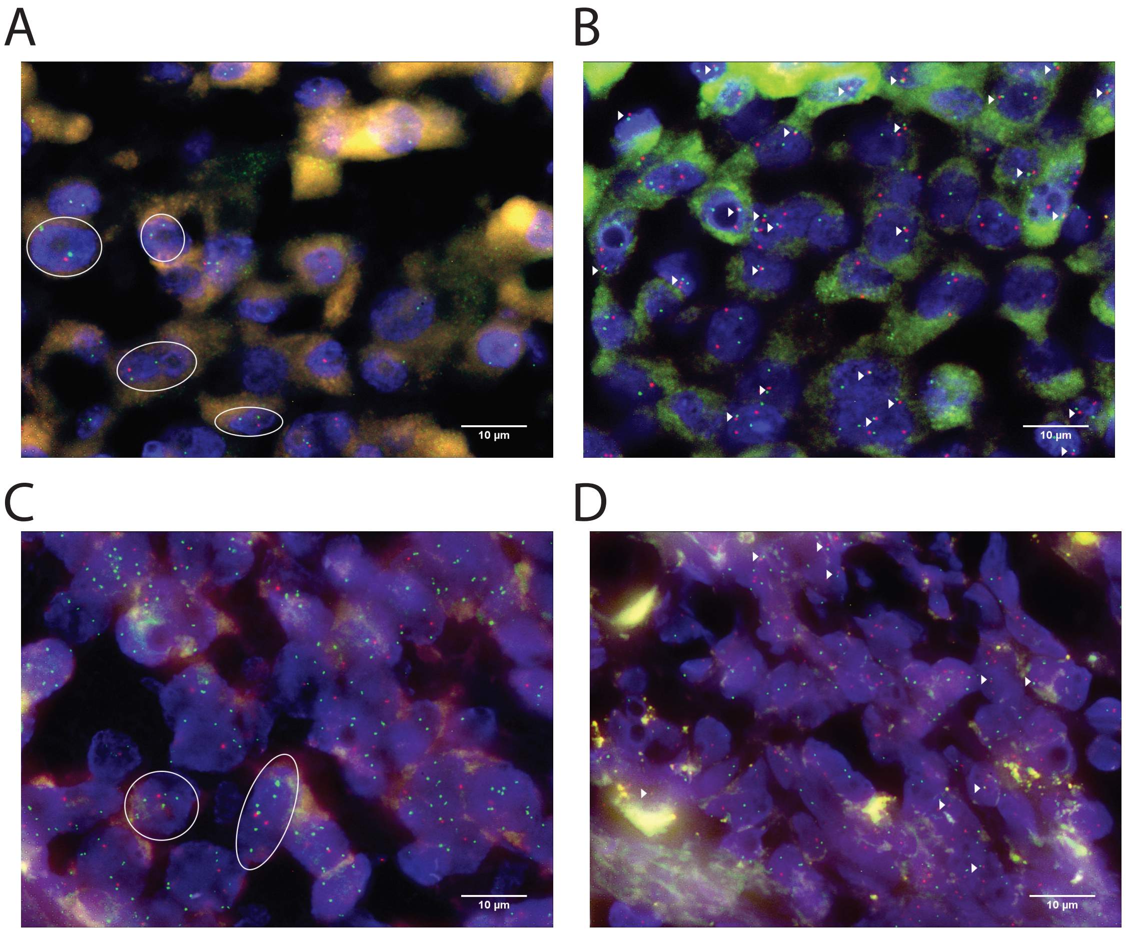


**Supplementary Figure 5. Two colour fluorescence in-situ hybridization (FISH) analyses confirm driver reorganizations in HCC tumours. (a)** FISH analysis of the *TP53* deletion in HCC SA529830. Representative cells harbouring the deletion are shown inside the circles. Two green signals (control probe) and one red signal (*TP53* probe) are detected in each cell. **(b)** Single-fusion FISH analysis of the t(8,17) translocation from the same tumour. In cells with the chromosomal fusion, one green and one red signal, each one targeting one the two chromosomes involved, colocalize together. In case of the standard (i.e., non-rearranged) karyotype, green and red signals split. Arrowheads point to the relevant chromosomal fusions. **(c)** FISH analysis of the *ARID1A* deletion in HCC SA501424. Representative cells harbouring the deletion are shown inside the circles.

Green signals (control probe) and red signals (*ARID1A* probe) are shown. In this case, between six to ten green signals and two red signals are detected in each cell, compatible with *ARID1A* deletion (see **Methods**). **(d)** Single-fusion FISH analysis of the t(1,11) translocation from the tumour SA501424. Arrowheads point to the relevant chromosomal fusions.

#### SUPPLEMENTARY TABLES

**Supplementary Table 1.** HBV insertions in human HCCs from the PCAWG dataset

**Supplementary Table 2.** Clonal state of HBV insertions

**Supplementary Table 3.** Real-time timing of whole genome duplication events along with HBV insertions clonal states
